# Disentangling genetically confounded polygenic associations between Attention-Deficit/Hyperactivity Disorder, literacy and language

**DOI:** 10.1101/276527

**Authors:** Ellen Verhoef, Ditte Demontis, Stephen Burgess, Chin Yang Shapland, Philip S. Dale, Aysu Okbay, Benjamin M. Neale, Stephen V. Faraone, iPSYCH-Broad-PGC ADHD Consortium, Evie Stergiakouli, George Davey Smith, Simon E. Fisher, Anders D. Børglum, Beate St Pourcain

## Abstract

Interpreting polygenic overlap between ADHD and both literacy- and language-related impairments is challenging as genetic confounding can bias associations. Here, we investigate evidence for links between polygenic ADHD risk and multiple literacy- and language-related abilities (LRAs), assessed in UK children (N≤5,919), conditional on genetic effects shared with educational attainment (EA). Genome-wide summary statistics on clinical ADHD and years-of-schooling were obtained from large consortia (N≤326,041). ADHD-polygenic scores (ADHD-PGS) were inversely associated with LRAs in ALSPAC, most consistently with reading-related abilities, and explained ≤1.6% phenotypic variation. Polygenic links were then dissected into both genetic effects shared with and independent of EA using multivariable regressions (MVR), analogous to Mendelian Randomization approaches accounting for mediating effects. Conditional on EA, polygenic ADHD risk remained associated with multiple literacy-related skills, phonemic awareness and verbal intelligence, but not language-related skills such as listening comprehension and non-word repetition. Pooled reading performance showed the strongest overlap with ADHD independent of EA. Using conservative ADHD-instruments (*P*-threshold<5×10^−8^) this corresponded to a 0.35 decrease in Z-scores per log-odds in ADHD-liability (*P*=9.2×10^−5^). Using subthreshold ADHD-instruments (*P*-threshold<0.0015), these associations had lower magnitude, but higher predictive accuracy, with a 0.03 decrease in Z-scores (*P*=1.4×10^−6^). Polygenic ADHD-effects shared with EA were of equal strength and at least equal magnitude compared to those independent of EA, for all LRAs studied, and only detectable using subthreshold instruments. Thus, ADHD-related polygenic links are highly susceptible to genetic confounding, concealing an ADHD-specific association profile that primarily involves reading-related impairments, but few language-related problems.

## Introduction

Children with Attention-Deficit/Hyperactivity Disorder (ADHD) often experience difficulties mastering literacy- and language-related abilities (LRAs)^1–3^. It has been estimated that up to 40% of children diagnosed with clinical ADHD also suffer from reading disability (RD, also known as developmental dyslexia) and vice versa^4^. The spectrum of affected LRAs in ADHD may, however, also include writing^5,6^, spelling^7,8^, syntactic^9,10^ and phonological^9,10^ abilities. Both clinical ADHD and RD are complex childhood-onset neurodevelopmental conditions that affect about 5% and 7% of the general population respectively^11,12^. ADHD is characterised by hyperactive, inattentive and impulsive symptoms^13^, whereas decoding and/or reading comprehension deficits are prominent in individuals with RD^14^.

To interpret the comorbidity of ADHD and RD a multiple-deficit model including shared underlying aetiologies has been proposed, involving both genetic and environmental influences^15^. This model is supported by twin studies suggesting that the co-occurrence of ADHD symptoms and reading deficits is to a large extent attributable to shared genetic influences^16–18^. However, the interpretation of polygenic overlap is challenging as associations might be inflated or induced by genetic confounders that are genetically related to both ADHD and reading abilities. Such a potential confounder of polygenic ADHD links is genetically predicted educational attainment (EA), a proxy of cognitive abilities and socioeconomic status^19^. Previous research showed evidence for a positive genetic relation between EA and reading abilities^20^, as well as a moderate negative genetic correlation between EA and ADHD^21^. Therefore, genetic effects shared with EA might obscure the genetic relationship between ADHD and LRAs. Both the extent and nature of the genetic overlap between ADHD and LRAs, accounting for genetic confounding through EA, are unknown.

Here, we (1) study polygenic links between clinical ADHD and a wide range of population-ascertained literacy- and language-related measures, (2) evaluate whether such links reflect a shared genetic basis with EA and (3) assess whether there is support for shared genetic factors between clinical ADHD and LRAs conditional on genetically predicted EA. ADHD polygenic scores (ADHD-PGS) are generated based on ADHD genome-wide association study (GWAS) summary statistics of two large independent ADHD samples, the Psychiatric Genomics Consortium (PGC) and the Danish Lundbeck Foundation Initiative for Integrative Psychiatric Research (iPSYCH) and a combination thereof. Associations between ADHD-PGS and a spectrum of population-based literacy- and language-related measures related to reading, spelling, phonemic awareness, listening comprehension, non-word repetition and verbal intelligence skills, are examined in a sample of children from the UK Avon Longitudinal Study of Parents and Children (ALSPAC). Applying multivariable regression (MVR) techniques, analogous to Mendelian Randomization (MR) approaches accounting for mediating effects^22^, we disentangled associations between polygenic ADHD risk and LRA measures into effects independent of and shared with years-of-schooling, using summary statistics from the Social Science Genetic Association Consortium (SSGAC).

## Methods and Materials

### Literacy- and language-related abilities in the general population

LRAs were assessed in children and adolescents from ALSPAC, a UK population-based longitudinal pregnancy-ascertained birth cohort (estimated birth date: 1991-1992, Supplementary Information)^23,24^. Ethical approval was obtained from the ALSPAC Law-and-Ethics Committee (IRB00003312) and the Local Research-Ethics Committees. Written informed consent was obtained from a parent or individual with parental responsibility and assent (and for older children consent) was obtained from the child participants.

#### Phenotype information

Thirteen measures capturing LRAs related to reading, spelling, phonemic awareness, listening comprehension, non-word repetition and verbal intelligence scores were assessed in 7 to 13 year-old ALSPAC participants (N≤5,919, Table 1) using both standardised and ALSPAC-specific instruments (Supplementary Information). Reading measures included comprehension and decoding (based on words and non-words) scores. Detailed descriptions of all LRA measures are available in Table 1 and the Supplementary Information.

**Table 1:**
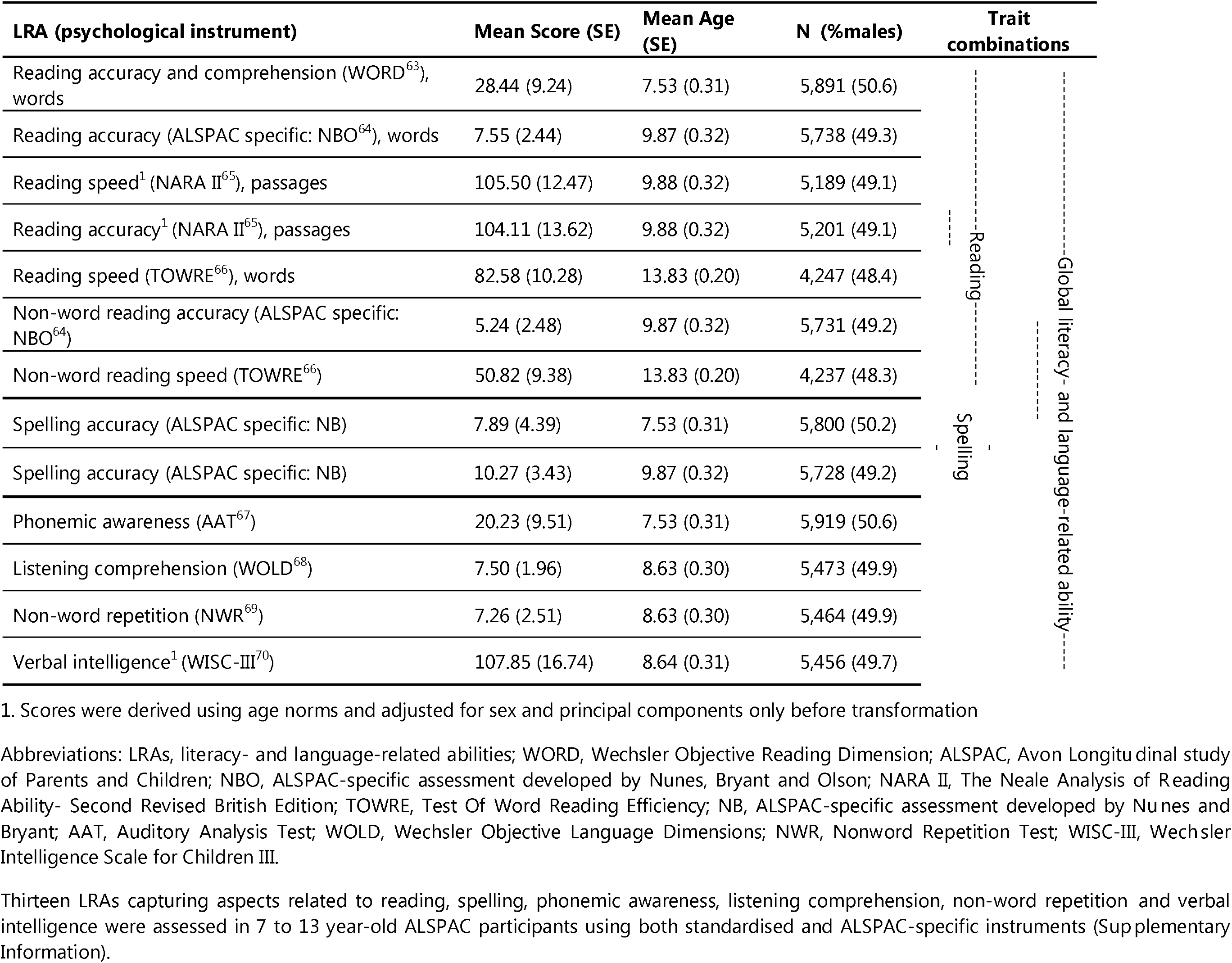
Literacy and literacy- and language-related oral language abilities in the Avon Longitudinal Study of Parents and Children

All LRA scores were rank-transformed to allow for comparisons of genetic effects across different psychological instruments with different distributions (Supplementary Information). Phenotypic correlations, using Pearson-correlation coefficients, were comparable for untransformed and rank-transformed scores (Table SI). To account for multiple testing, we estimated the effective number of phenotypes studied using Matrix Spectral Decomposition^25^ (MatSpD), revealing seven independent measures (experiment-wide error rate of 0.007).

For sensitivity analysis, we excluded 188 children with LRA data available and an ADHD diagnosis at age 7, based on the Development and Wellbeing Assessment (DAWBA)^26^ (Supplementary Information).

#### Genetic analyses

ALSPAC participants were genotyped using the Illumina HumanHap550 quad chip genotyping platforms, and genotypes were called using the Illumina GenomeStudio software. Genotyping, imputation and genome-wide association analysis details are described in the Supplementary Information and Table 2.

**Table 2:**
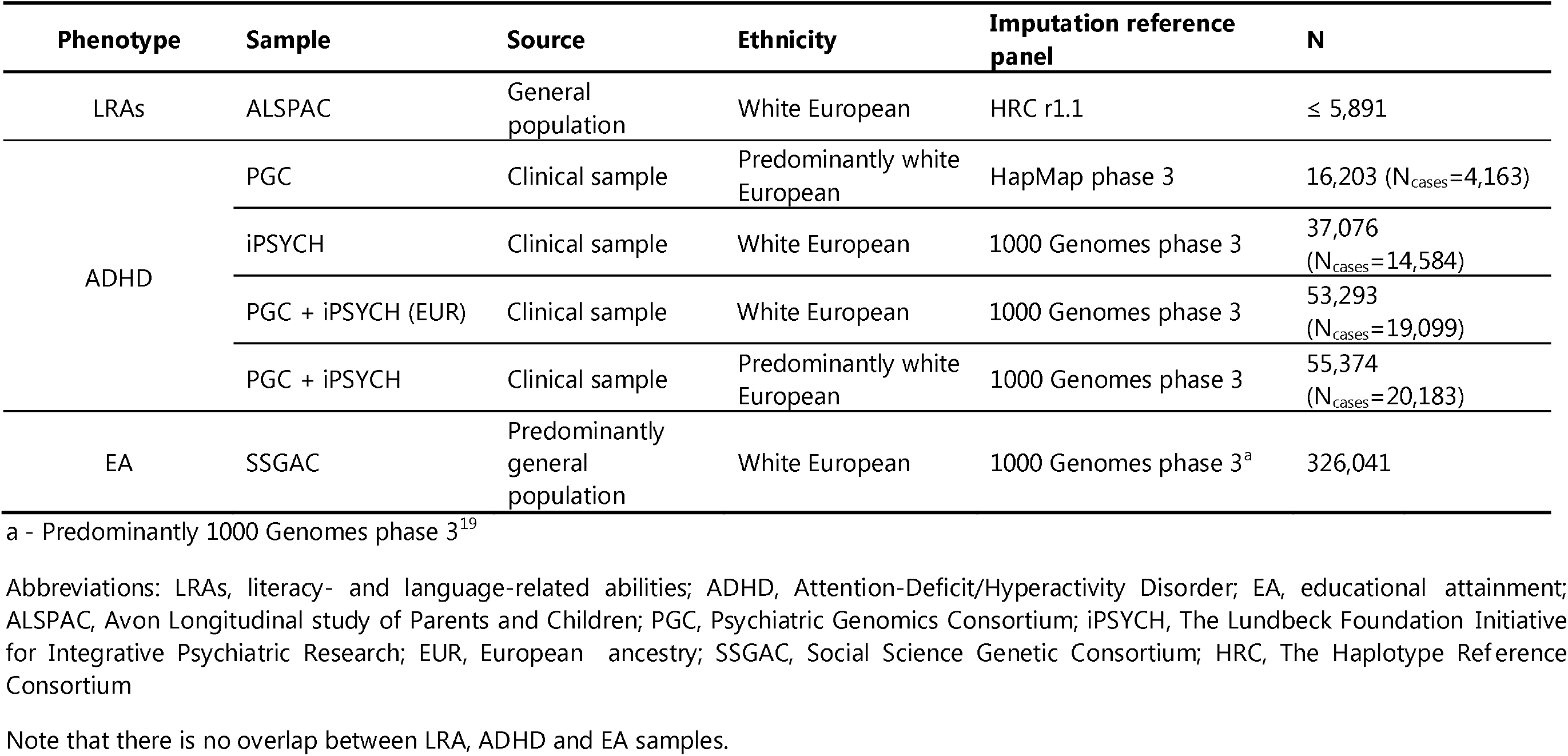
Sample description

### Clinical ADHD summary statistics

#### Psychiatric Genomics Consortium (PGC)

GWAS summary statistics were obtained from a mega-analysis of clinical ADHD^27^, conducted by the PGC (4,163 cases and 12,040 controls/pseudo-controls) (Table 2, Supplementary Information, www.med.unc.edu/pgc/).

#### The Lundbeck Foundation Initiative for Integrative Psychiatric Research (iPSYCH)

An independent set of ADHD GWAS summary statistics were accessed through the Danish iPSYCH project^28^ (14,584 ADHD cases, 22,492 controls) (Table 2, Supplementary Information), using samples from the Danish Neonatal Screening Biobank hosted by Statens Serum Institute^21,29^.

#### Combined PGC and iPSYCH ADHD sample (PGC+iPSYCH)

To maximise power, we also analysed meta-GWAS summary statistics from an ADHD sample containing both PGC and iPSYCH participants^21^ (20,183 cases, 35,191 controls/pseudo-controls) (Table 2, www.med.unc.edu/pgc/) and its European-only subset (*PGC+iPSYCH(EUR*), 19,099 cases, 34,194 controls/pseudo-controls) (Table 2, www.med.unc.edu/pac/).

Detailed sample descriptions are available in Table 2 and the Supplementary Information.

### Educational attainment summary statistics

GWAS summary statistics for EA^19^ (discovery and replication sample combined, excluding ALSPAC and 23andMe samples, N=326,041) were obtained from the SSGAC consortium. EA was assessed as years of schooling^19^. A detailed sample description is available in Table 2 and the Supplementary Information.

### Genome-wide complex trait analysis

SNP-h^2^ and genetic correlations (r_g_) between LRAs were estimated using Restricted Maximum Likelihood (REML) analyses^30,31^ as implemented in Genome-wide Complex Trait Analysis (GCTA) software^32^, including individuals with a genetic relationship <0.05^30^. For this study, we selected only LRAs with evidence for SNP-h^2^ and sample size N>4,000 (Table S2).

### Linkage Disequilibrium Score regression and correlation

Linkage Disequilibrium Score (LDSC) regression^33^ was used to distinguish confounding biases from polygenic influences by examining the LDSC regression intercept. Unconstrained LD-score correlation^34^ analysis was applied to estimate r_g_ (Supplementary Information).

### Polygenic scoring analyses

ADHD-PGS^35,36^ were created in ALSPAC using PGC, iPSYCH and PGC+iPSYCH GWAS summary statistics (Supplementary Information). ADHD-PGS have been previously linked to ADHD symptoms in ALSPAC participants^37^. Rank-transformed LRAs were regressed on Z-standardised ADHD-PGS (aligned to measure risk-increasing alleles) using ordinary least square (OLS) regression (R:stats library, Rv3.2.0). The proportion of phenotypic variance explained is reported as OLS-regression-R^2^. Beta-coefficients (β) for ADHD-PGS quantify here the change in standard deviation (SD) units of LRA performance per one SD increase in ADHD-PGS.

### Multivariable regression analysis

To study the genetic association between ADHD and LRAs conditional on genetic influences shared with EA, we applied MVR. This technique is analogous to MR methodologies accounting for mediating effects^22^ and controls for collider bias^38^ through the use of GWAS summary statistics. Technically, it involves the regression of regression estimates from independent samples on each other^22^ (Supplementary Information). Within this study we use MVR without inferring causality due to violations of classical MR assumptions^22^ (see below).

#### Genetic variant selection

To disentangle ADHD-LRA associations, we selected two sets of genetic variants from the most powerful ADHD GWAS summary statistics (PGC+iPSYCH). The first set contained genome-wide significant variants (*P*<5×10^−8^, conservative). The second set included variants passing a more lenient *P*-value threshold (*P*<0.0015, subthreshold) to increase power, consistent with current guidelines for the selection of genetic instruments in MR (F-statistic<10)^39^. All sets included independent (PLINK^40^ clumping: LD-r^2^<0.25, ±500 kb), well imputed (INFO^41^>0.8) and common (EAF>0.01) variants. This resulted in 15 conservative and 2,689<N_SNPs_≤2,692 subthreshold ADHD-instruments (Table S8).

#### Estimation of ADHD effects

We extracted regression estimates for selected ADHD-instruments (conservative and subthreshold) from ADHD (PGC+iPSYCH), EA (SSGAC) and LRA (ALSPAC) GWAS summary statistics. Analysing each set of variants independently, regression estimates for individual LRA measures (β) were regressed on both ADHD (β as lnOR) and EA regression estimates (β) using an OLS regression framework (R:stats library, Rv3.2.0). Outcomes were 1) a MVR regression estimate quantifying the change in SD units of LRA performance per log odds increase in ADHD risk conditional on years of schooling (ADHD effect independent of EA), and 2) a MVR regression estimate quantifying the change in SD units of LRA performance per year of schooling as captured by ADHD instruments (ADHD effect shared with EA). Latter MVR regression estimates capture here genetic confounding, including confounding influences in a narrow sense (i.e. genetically predictable EA causally influences both ADHD and LRA), but also mediating effects (i.e. genetically predictable ADHD causally influences LRA indirectly through EA) and biological pleiotropy (i.e. ADHD risk variants affect ADHD and EA through independent biological pathways). As ADHD risk and EA are inversely genetically related with each other^21^, they were converted to quantify a change per missing year of schooling. To compare the magnitude of both MVR estimates, we also conducted analyses using fully standardised EA, ADHD and LRA regression estimates (Supplementary Information).

Finally, MVR regression estimates were meta-analysed across reading-related, spelling-related and all LRA measures (Table 1) using random-effects meta-regression, accounting for phenotypic correlations between LRAs (R:metafor library, Rv3.2.0, Supplementary Information).

#### Sensitivity analyses

As the directionality of effects cannot be inferred in this study, we examined, in reverse, evidence for ADHD as a genetic confounder of the association between EA and LRAs, using MVR. Two sets of EA instruments (conservative and subthreshold, Table S8) were selected from EA (SSGAC) GWAS summary statistics, analogous to the selection of ADHD instruments, and MVR was conducted as described above.

### Attrition analysis

We carried out an *attrition* analysis in ALSPAC studying the genetic association between LRA-missingness and ADHD-PGS (Supplementary Information).

## Results

### Genetic architecture of literacy- and lanαuaαe-related abilities and clinical ADHD

Phenotypic variation in literacy- and language-related measures (Table 1), including reading abilities (comprehension, accuracy and speed) assessed in words/passages and non-words, spelling abilities (accuracy), phonemic awareness, listening comprehension, non-word repetition and verbal intelligence scores, can be tagged by common variants, with SNP-h^2^ estimates between 0.32 (SE=0.07, non-word repetition age 8) and 0.54 (SE=0.07, verbal intelligence age 8) (Table S2; GCTA- and LDSC-based estimations). All LRAs are phenotypically (Table SI) and genetically (Table S3) moderately to strongly interrelated. The observed LDSC-based evidence for genetic liability of clinical ADHD within the PGC (LDSC-h^2^=0.08(SE=0.03)), iPSYCH (LDSC-h^2^=0.26(SE=0.02)) and PGC+iPSYCH samples (Table S4) is consistent with previous reports^21^.

### Association between ADHD polygenic risk scores and literacy- and language-related abilities

We observed robust evidence for an inverse genetic association between ADHD-PGS and reading accuracy/comprehension age 7 (PGC: OLS-R^2^=0.1%, *P*=4.6×10^−3^; iPSYCH: OLS-R^2^=1.0%, *P*<1×^−10^), reading accuracy age 9 (PGC: OLS-R^2^=0.1%, *P*=5.7×10^−3^; iPSYCH: OLS-R^2^=1.2%, *P*<1×^−10^), and spelling accuracy age 9 (PGC: OLS-R^2^=0.2%, *P*=1.5×10^−3^; iPSYCH: OLS-R^2^=C>.8%, *P*<1×10^−10^) using independent ADHD discovery samples (Figure 1, Table S5). Polygenic scoring results are presented for a P-value threshold of 0.1, but other thresholds provided similar results (data not shown). The strongest evidence for association was observed when ADHD discovery samples were combined (PGC+iPSYCH; Figure 1, Table S5), including those of European ancestry only (PGC+iPSYCH(EUR), Table S5), and genetic overlap was present for all LRAs studied. Results were not affected by the exclusion of children with an ADHD diagnosis in ALSPAC (Table S6).

**Figure 1:**
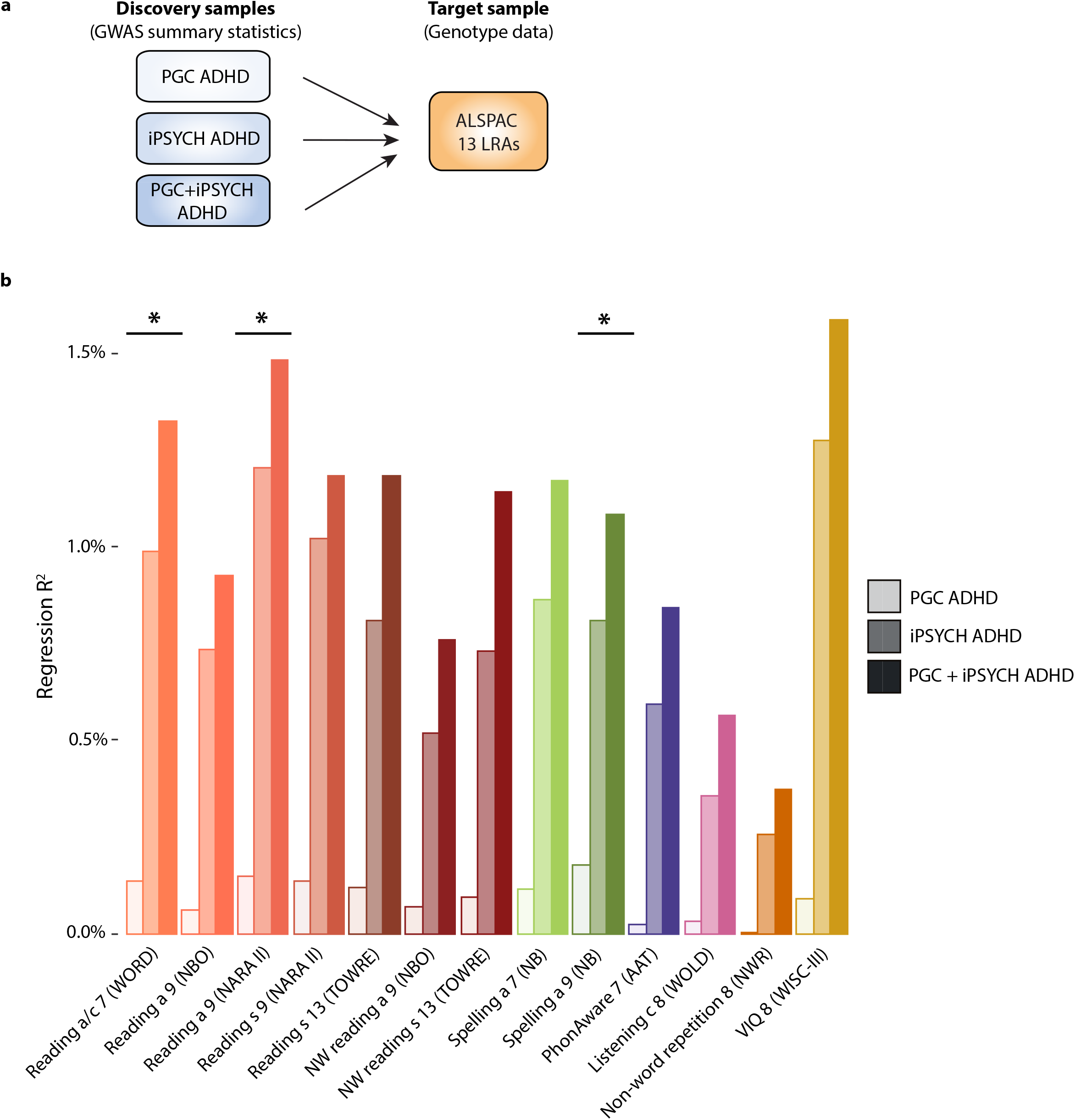
Phenotypic variance in literacy- and language-related abilities explained by polygenic ADHD risk. Abbreviations: a, accuracy; c, comprehension; s, speed; WORD, Wechsler Objective Reading Dimension; NBO, Nunes, Bryant and Olson (ALSPAC specific instrument); NARA II, The Neale Analysis of Reading Ability-Second Revised British Edition; TOWRE, Test Of Word Reading Efficiency; NW, non-word; NB, Nunes and Bryant (ALSPAC specific instrument); PhonAware, phonemic awareness; AAT, Auditory Analysis Test; WOLD, Wechsler Objective Language Dimensions; NWR, Non-word Repetition Test; VIQ, verbal intelligence quotient; WISC-III, Wechsler Intelligence Scale for Children III; PGC, Psychiatric Genomics Consortium; iPSYCH, The Lundbeck Foundation Initiative for Integrative Psychiatric Research; ADHD, Attention-Deficit/Hyperactivity Disorder. a) Schematic representation of polygenic scoring analyses. ADHD polygenic scores were created in ALSPAC using PGC, iPSYCH and PGC+iPSYCH GWAS summary statistics. Rank-transformed LRAs were regressed on Z-standardised ADHD-PGS using ordinary least square regression, b) Phenotypic variance in literacy- and language-related abilities explained by polygenic ADHD risk. *-Evidence for association between LRAs and polygenic ADHD risk as observed in PGC ADHD, iPSYCH ADHD and PGC+iPSYCH ADHD samples. Note that all LRAs were associated with polygenic ADHD risk in iPSYCH ADHD and PGC+iPSYCH ADHD passing the experiment-wide error rate (*P*<0.007).

### Shared genetic liability between ADHD and LRA with EA

There was strong evidence for a moderate negative genetic correlation (r_g_=−0.53(SE=0.03), *P*<1×10^−10^) between genetically predicted ADHD, as captured by the largest ADHD discovery sample (PGC+iPSYCH), and EA (LDSC-h^2^=0.11(SE=0.004)), consistent with previous findings^21^. Likewise, LRAs were moderately to highly positively correlated with EA (e.g. reading speed age 13 r_g_=0.80(SE=0.22), *P*=3.0×10^−4^; Table S7), as previously reported^20^. Additionally, two independent variants reached genome-wide significance for both ADHD^21^ and EA^19^, consistent with biological pleiotropy^42^ (i.e. single genetic loci influencing multiple traits)^42^. These findings indicate complex, potentially reciprocal cross-trait relationships (Figure 2a) and violate MR causal modelling assumptions^22^. Consequently, ADHD instruments are not valid MR instruments as they are not independent of EA, a potential genetic confounder.

**Figure 2:**
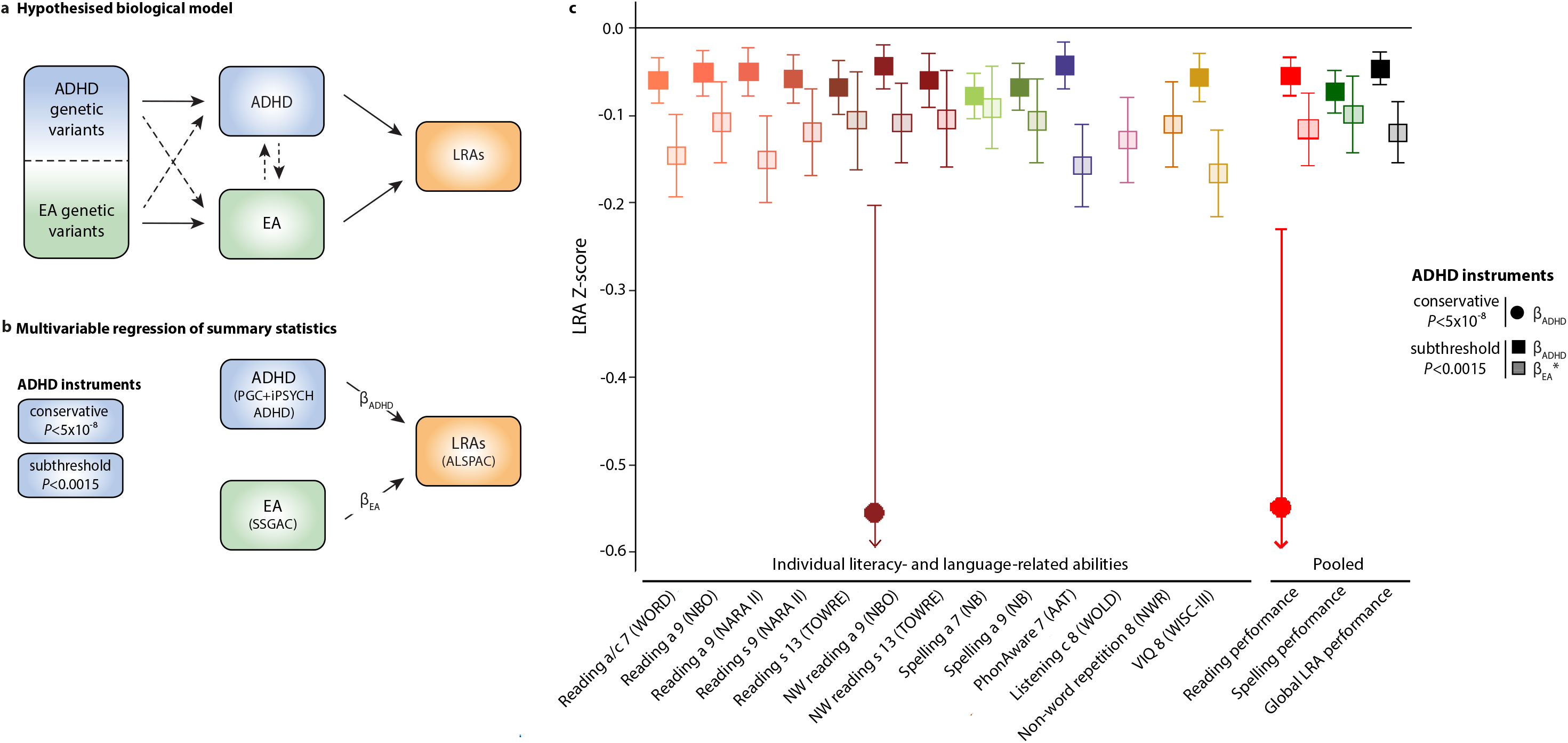
Genetic relationships between ADHD, educational attainment and literacy- and language-related abilities. Abbreviations: ADHD, Attention-Deficit/Hyperactivity Disorder; EA, educational attainment; LRAs, literacy and language-related abilities; PGC, Psychiatric Genomics Consortium; iPSYCH, The Lundbeck Foundation Initiative for Integrative Psychiatric Research; SSGAC, Science Genetic Association Consortium; ALSPAC, Avon Longitudinal Study of Parents And Children; MVR, multivariable regression (a) Hypothesized biological model of genetic relationships between ADHD, EA, and LRAs, including complex, pleiotropic and reciprocal genetic links, (b) Schematic MVR model assessing polygenic ADHD-LRA overlap independent of and shared with genetic effects for EA, (c) MVR estimates of ADHD-specific effects independent of EA and ADHD effects shared with EA on LRAs using standardised ADHD instruments: Sets of conservative (*P*<5×10^−8^) and subthreshold (*P*<0.0015) ADHD instruments were extracted from ADHD (PGC+iPSYCH), EA (SSGAC) and LRAs (ALSPAC) GWAS summary statistics. ADHD-specific effects independent of EA (PADHD) and ADHD effects shared with EA (PEA) on LRAs were estimated with MVRs. To compare the magnitude of Padhd and p_EA_, MVR analyses were conducted using standardised regression estimates (Supplementary Methods). PADHD estimates measure the change in LRA Z-score per Z-score in ADHD liability. Pea estimates measure the change in LRA Z-scores per Z-score in missing school years. MVR estimates based on raw genetic effect estimates are provided in Table 3. Pooled estimates for reading, spelling and global LRA measures (Table 1) were obtained through random-effects meta-regression. Only effects passing the experiment-wide significance threshold (*P*<0.007) are shown with corresponding 95% confidence intervals. There is no causality inferred. * ADHD effects shared with EA were assessed through EA genetic effect estimates of ADHD-associated variants.

### Multivariable regression analyses

To disentangle the genetic overlap of polygenic ADHD risk with literacy- and language-related measures into genetic ADHD effects independent of and shared with EA, we applied MVR^22^ using ADHD instruments based on the most powerful ADHD discovery sample (PGC+iPSYCH) (Figure 2b).

Using conservative ADHD instruments (Table S8), only non-word reading accuracy age 9 was associated with polygenic ADHD conditional on EA, with a decrease of 0.37 SD per log-odds increase in ADHD risk (β_ADHD_=−0.37(SE=0.09), *P*=1.7×10^−3^) (Table 3). Additionally, when meta-analysing LRAs across pre-defined domains, we observed an effect of polygenic ADHD risk on pooled reading performance, independent of EA. This corresponds to a decrease of 0.35 SD in pooled reading performance per log-odds increase in ADHD liability (β_ADHD_=−0.35(SE=0.09), *P*=9.2×10^−5^, *P_het_*=0.19). There was little evidence for association with spelling or combined LRA performance conditional on EA (Table 3).

**Table 3:** Multivariable regression analysis of polygenic associations between ADHD and literacy- and language-related abilities (raw estimates)

| LRAs | ADHD ($\beta_{\text{ADHD}}$)<br>(ADHD-specific effects independent of EA) | | | | | | EA ($\beta_{\text{EA}}$)<br>(EA genetic effects of ADHD-associated variants)<sup>1</sup> | | | | | |
| --- | --- | --- | --- | --- | --- | --- | --- | --- | --- | --- | --- | --- |
| | Conservative instruments<br>($P_{\text{thr}} < 5 \times 10^{-8}$) | | | Subthreshold<br>instruments ($P_{\text{thr}} < 0.0015$) | | | Conservative instruments<br>($P_{\text{thr}} < 5 \times 10^{-8}$) | | | Subthreshold instruments<br>($P_{\text{thr}} < 0.0015$) | | |
| | $\beta$ (SE) | $P$ | $P_{\text{het}}$ | $\beta$ (SE) | $P$ | $P_{\text{het}}$ | $\beta$ (SE) | $P$ | $P_{\text{het}}$ | $\beta$ (SE) | $P$ | $P_{\text{het}}$ |
| Reading a/c 7 (WORD) | -0.18<br>(0.13) | 0.20 | - | -0.03<br>(0.01) | $1.4 \times 10^{-5}$ | - | -0.09<br>(1.19) | 0.94 | - | -0.63(0.10) | $1.2 \times 10^{-9}$ | - |
| Reading a 9 (NBO) | -0.28<br>(0.09) | 0.01 | - | -0.03<br>(0.01) | $2.1 \times 10^{-4}$ | - | 1.33 (0.84) | 0.14 | - | -0.46 (0.10) | $6.3 \times 10^{-6}$ | - |
| Reading s 9 (NARA II) | -0.29<br>(0.13) | 0.04 | - | -0.03<br>(0.01) | $3.0 \times 10^{-5}$ | - | 1.55 (1.16) | 0.21 | - | -0.51 (0.11) | $2.5 \times 10^{-6}$ | - |
| Reading a 9 (NARA II) | -0.34<br>(0.12) | 0.01 | - | -0.03<br>(0.01) | $3.8 \times 10^{-4}$ | - | 1.57 (1.11) | 0.18 | - | -0.64 (0.11) | $4.1 \times 10^{-9}$ | - |
| Reading s 13 (TOWRE) | -0.36<br>(0.15) | 0.03 | - | -0.04 (0.01) | $2.7 \times 10^{-5}$ | - | 2.06 (1.35) | 0.15 | - | -0.46 (0.12) | $1.8 \times 10^{-4}$ | - |
| NW reading a 9 (NBO) | -0.37<br>(0.09) | $1.7 \times 10^{-3}$ | - | -0.03<br>(0.01) | $2.4 \times 10^{-4}$ | - | 1.54 (0.85) | 0.09 | - | -0.46 (0.10) | $6.1 \times 10^{-6}$ | - |
| NW reading s 13 (TOWRE) | -0.21<br>(0.17) | 0.25 | - | -0.03<br>(0.01) | $2.4 \times 10^{-4}$ | - | 1.43 (1.58) | 0.38 | - | -0.45 (0.12) | $1.9 \times 10^{-4}$ | - |
| Spelling a 7 (NB) | -0.05<br>(0.12) | 0.72 | - | -0.05<br>(0.01) | $1.4 \times 10^{-5}$ | - | -1.12<br>(1.13) | 0.34 | - | -0.63 (0.10) | $1.2 \times 10^{-9}$ | - |
| Spelling a 9 (NB) | -0.23<br>(0.08) | 0.01 | - | -0.04<br>(0.01) | $8.7 \times 10^{-7}$ | - | 1.43 (0.72) | 0.38 | - | -0.45 (0.10) | $1.3 \times 10^{-5}$ | - |
| PhonAware 7 (AAT) | -0.22<br>(0.15) | 0.16 | - | -0.02<br>(0.01) | $1.9 \times 10^{-3}$ | - | -0.31<br>(1.33) | 0.82 | - | -0.67 (0.10) | $< 1 \times 10^{-10}$ | - |
| Listencing c 8 (WOLD) | -0.03<br>(0.08) | 0.75 | - | -0.02<br>(0.01) | 0.02 | - | -1.47<br>(0.70) | 0.06 | - | -0.55 (0.11) | $4.5 \times 10^{-7}$ | - |
| Non-word repetition 8 (NWR) | -0.14<br>(0.14) | 0.34 | - | -0.02<br>(0.01) | 0.01 | - | -0.16<br>(1.30) | 0.90 | - | -0.47 (0.11) | $1.5 \times 10^{-5}$ | - |
| VIQ 8 (WISC-III) | -0.23<br>(0.15) | 0.16 | - | -0.03<br>(0.01) | $5.0 \times 10^{-5}$ | - | -0.36<br>(1.37) | 0.80 | - | -0.71 (0.11) | $< 1 \times 10^{-10}$ | - |
| Pooled reading performance | -0.35<br>(0.09) | $9.2 \times 10^{-5}$ | 0.19 | -0.03<br>(0.01) | $1.4 \times 10^{-6}$ | 0.79 | 1.67 (0.78) | 0.03 | 0.31 | -0.50 (0.09) | $4.9 \times 10^{-8}$ | 0.09 |
| Pooled spelling performance | -0.18<br>(0.11) | 0.10 | 0.03 | -0.04<br>(0.01) | $1.1 \times 10^{-8}$ | 0.28 | 0.40 (1.25) | 0.75 | 0.001 | -0.42 (0.10) | $1.3 \times 10^{-5}$ | 0.39 |
| Global LRA performance | -0.18<br>(0.07) | 0.01 | 0.007 | -0.03<br>(0.01) | $1.2 \times 10^{-6}$ | 0.06 | 0.29 (0.68) | 0.67 | 0.002 | 0.51 (0.08) | $< 1 \times 10^{-10}$ | 0.002 |

1. ADHD genetic effects shared with EA as assessed through EA genetic effect estimates of ADHD-associated variants
Abbreviations: LRAs, literacy- and language-related abilities; ADHD, Attention-Deficit/Hyperactivity Disorder; EA, educational attainment; $P_{thr}$ , $P$ -value threshold; $P$ , $P$ -value; $P_{het}$ , heterogeneity $P$ -value; a, accuracy; c, comprehension; s, speed; WORD, Wechsler Objective Reading Dimension; NBO, Nunes, Bryant and Olson (ALSPAC specific instrument); NARA II, The Neale Analysis of Reading Ability- Second Revised British Edition; TOWRE, Test Of Word Reading Efficiency; NW, non-word; NB, Nunes and Bryant (ALSPAC specific instrument); PhonAware, phonemic awareness; AAT, Auditory Analysis Test; WOLD, Wechsler Objective Language Dimensions; NWR, Nonword Repetition Test; VIQ, verbal intelligence quotient; WISC-III, Wechsler Intelligence Scale for Children III; MVR - Multivariable regression.

Using subthreshold ADHD instruments (Table S8), polygenic ADHD effects on LRA performance, conditional on EA, were detectable for all reading- and spelling-related measures, phonemic awareness and verbal intelligence, but not language-related skills such as listening comprehension and non-word repetition (Table 3). Observable effects were, however, considerably smaller than those captured by conservative ADHD instruments although the statistical evidence increased, such as a 0.03 SD decrease in non-word reading accuracy at age 9 per log-odds increase in ADHD risk (β_ADHD_=−0.03(SE=0.01), *P*=2.4×10^−4^, Table 3).

Polygenic ADHD effects on literacy- and language-related performance that are shared with EA were identified for all LRAs studied, but only using subthreshold, not conservative ADHD instruments (Table 3). This translates into, for example, a further 0.46 SD units decrease in non-word reading accuracy at age 9 per missing school year (β_EA_=−0.46(SE=0.10), *P*=6.1×10^−6^, Table 3). Thus, the association between polygenic ADHD risk and listening comprehension and non-word repetition, is fully attributable to genetic effects shared with EA (Table 3). MVR using fully standardised EA, ADHD and LRA regression estimates showed that ADHD effects shared with EA were as large as or even larger than ADHD effects independent of EA (Figure 2c, Table S9).

Using an analogous approach, we also disentangled the genetic overlap of polygenic EA with LRAs into genetic EA effects independent of and shared with ADHD based on EA instruments (Figure S1). As observed using ADHD instruments, there was strong evidence for EA effects shared with ADHD using subthreshold, but not conservative EA instruments (Table S10). Importantly, the magnitude of ADHD genetic effects shared with EA compared to EA genetic effects shared with ADHD was largely consistent with each other in fully standardised analyses (Table S9 and S10).

All MVRs were performed using constrained intercepts^22^. There was little evidence supporting the inclusion of regression intercepts that would imply additional genetic effects using these instruments, not yet captured by either ADHD or EA effect estimates.

### Attrition in ALSPAC

Analyses of sample drop-out in ALSPAC, exemplified by missing reading accuracy and comprehension scores at age 7 (WORD), were carried out to investigate potential sources of bias. We identified a positive genetic association between sample-dropout and ADHD-PGS (PGC+iPSYCH, OR=1.03(SE=0.005), *P*=1.4×10^−8^, Table S11), consistent with previous studies^43^. Disentangling this polygenic link using MVR (Table S12) based on ADHD subthreshold instruments showed a 0.05 increase in log-odds sample drop-out per log-odds increase in liability to ADHD independent of EA (log(OR)=0.05(SE=0.01), *P*=3.7×10^−4^). Additionally, a further 1.04 increase in log odds sample drop-out per missing year of schooling (log(OR)=1.04(SE=0.19),*P*=7.3×^−8^) was observed.

## Discussion

This study identified strong evidence for an inverse association between polygenic ADHD risk and multiple population-based LRAs using a polygenic scoring approach. However, these associations are genetically confounded through effects shared with genetically predictable EA. Accurate modelling of polygenic links using MVR techniques, conditional on EA, revealed an ADHD-specific association profile that primarily involves literacy-related impairments, but few language-related problems. Once genetic confounding is accounted for, polygenic ADHD risk was most strongly inversely associated with an overarching reading domain, in addition to spelling, phonemic awareness and verbal intelligence, but not listening comprehension and non-word repetition abilities. Importantly, ADHD genetic effects shared with EA inflated or even induced the genetic overlap between polygenic ADHD risk and all LRAs studied.

These findings are consistent with previous clinical and twin studies reporting a shared underlying aetiology between ADHD and literacy-related impairments^1–3,6–8,10,44,45^. They also reflect the observed robust evidence for an association between ADHD-PGS and several reading- and spelling-related abilities, using independent ADHD discovery samples.

The identified association profile suggests that reading-related deficits in ADHD may, as hypothesised for RD, arise from a phonological impairment^46,47^, which affects decoding and reading skills^48^, but also spelling abilities^49^. However, reading abilities can, once developed, also shape phonological skills^50^.

In addition, this study supports the genetic confounding of associations between polygenic ADHD and LRAs through genetically predictable EA, and, equally likely, the genetic confounding of associations between polygenic EA and LRAs through genetically predictable ADHD. The magnitude of these confounding effects was comparable, consistent with a model of reciprocal genetic influences between EA and ADHD (Figure 2a). Importantly, both directions of effect are supported by observational^51^ and twin^52^ studies. Thus, more generally, our findings support a multiple-deficit model proposing shared underlying aetiologies between ADHD and RD dimensions^15^.

Disentangling the underlying aetiological relationships between ADHD, EA and LRAs is challenging due to violations of classical MR assumptions^22^. This study focusses therefore on dissecting the genetic association between ADHD and LRAs into ADHD genetic effects that are shared with or independent of genetic confounders, such as EA, while controlling for collider bias^38^. Using MVR, the detection of these polygenic associations was strongly governed by the choice of genetic variants. Conservative ADHD instruments identified large ADHD-specific effects on reading as a domain, independent of EA, and little evidence for genetic effects shared with EA. However, conservative instruments had limited power^53^. They comprised 15 independent SNPs only, but included genetic variation within *FOXP2*, a gene that has been implicated in childhood apraxia of speech as well as expressive and receptive language impairments (http://omim.org/entry/602081)^54^. Subthreshold instruments including thousands of variants, on the other hand, tagged ADHD-specific polygenic associations (conditional on EA) of smaller magnitude, but with higher predictive accuracy. Moreover, these instruments detected strong evidence for genetic confounding affecting all polygenic links between ADHD and LRAs studied, with associations of equal strength and at least equal magnitude compared to associations independent of EA. In general, these findings are thus in support of an omnigenic^55^ model of complex trait architectures, compatible with a general factor model of psychopathology^56^, including ADHD^57^. The omnigenic model construes that only the largest-effect variants will reflect ADHD specificity, and may thus tag the most trait-specific associations between ADHD and reading, independent of EA. The majority of variants, however, will capture pleiotropic (omnigenic) influences pointing to highly interconnected neural networks^55^ that give rise to genetic confounding. Consequently, the majority of subthreshold variants, captured by both ADHD and EA subthreshold variants, are likely to represent highly powerful cross-trait genetic predictors that enhance and induce genetic overlap.

Finally, the methodological framework within this study has relevance not only for studies investigating polygenic links between ADHD and LRAs, but all studies examining polygenic overlap between ADHD and phenotypes that share genetically liability with EA or any other source of genetic confounding. Specifically, our findings suggest that lower variant selection thresholds can introduce genetic confounding that needs to be accounted for before identified associations can be interpreted in terms of underlying mechanisms, including shared genetic aetiologies. This is especially important as current guidelines for studying polygenic links with allelic scores recommend aggregating genetic variants across less stringent significance thresholds to maximise genetic association between discovery and target samples^58,59^.

This study has several limitations. Firstly, ALSPAC, as other cohort studies, suffers from attrition^43,60^. Sensitivity analyses showed that this is unlikely to bias our findings based on conservative instruments. However, links identified using subthreshold ADHD variants, might have been underestimated given that individuals with a genetic predisposition to ADHD are more likely to drop out^43^. Secondly, the strength of the genetic overlap between polygenic ADHD risk and LRAs may vary according to ADHD symptom domain levels, implicating especially inattentiveness^61^, as well as the nature of the literacy- or language-related ability involved (as we observed evidence for effect heterogeneity when combining all LRAs). It is conceivable that also other verbal abilities, not investigated in this study, such as grammar, expressive vocabulary or pragmatic skills, may genetically overlap with ADHD. Finally, we only studied evidence for genetic confounding with respect to genetically predictable EA using ADHD instruments (and vice versa genetically predictable ADHD using EA instruments). However, we found little evidence for the presence of additional unknown confounders, not yet tagged by these instruments. Furthermore, the power of available LRA GWAS statistics is still too low to generate genetic instruments supporting reverse models. Larger and more detailed clinical and population-based samples, as well as extensive multivariate variance analyses of the spectrum of LRAs (that are currently computationally expensive^62^) will help to discover and characterize further overlap between ADHD and literacy- and language-related cognitive processes.

## Conclusion

Polygenic ADHD links with LRAs are susceptible to genetic confounding through EA, especially when investigated with genetic variants typically selected for polygenic scoring approaches, inflating and biasing genetic overlap. Accounting for these unspecific genetic effects, reveals an ADHD-specific association profile that primarily involves reading-related impairments, but few language-related problems.

## Acknowledgements

We are extremely grateful to all the families who took part in this study, the midwives for their help in recruiting them, and the whole ALSPAC team, which includes interviewers, computer and laboratory technicians, clerical workers, research scientists, volunteers, managers, receptionists and nurses. The UK Medical Research Council and the Wellcome Trust (Grant ref: 102215/2/13/2) and the University of Bristol provide core support for ALSPAC. The ALSPAC GWAS data was generated by Sample Logistics and Genotyping Facilities at the Wellcome Trust Sanger Institute and LabCorp (Laboratory Corporation of America) using support from 23andMe.

This research was facilitated by the Social Science Genetic Association Consortium (SSGAC) and Psychiatric Genomics Consortium (PGC) by providing access to genome-wide summary statistics.

This publication is the work of the authors and EV and BSTP will serve as guarantors for the contents of this paper. EV, CYS, BSTP and SEF are supported by the Max Planck Society. The iPSYCH project is funded by the Lundbeck Foundation (grant no R102-A9118 and R155-2014-1724) and the universities and university hospitals of Aarhus and Copenhagen. Genotyping of iPSYCH and PGC samples was supported by grants from the Lundbeck Foundation, the Stanley Foundation, the Simons Foundation (SFARI 311789 to Mark J Daly), and NIMH (5U01MH094432-02 to Mark J Daly). The Danish National Biobank resource was supported by the Novo Nordisk Foundation. Data handling and analysis was supported by NIMH (1U01MH109514-01 to Michael O’Donovan and Anders D Børglum). High-performance computer capacity for handling and statistical analysis of iPSYCH data on the GenomeDK HPC facility was provided by the Centre for Integrative Sequencing, iSEQ, Aarhus University, Denmark (grant to Anders D Børglum).

An earlier version of this manuscript is available as a preprint on bioRxiv.

